# Critical role for Slam/SAP signaling in the thymic developmental programming of IL-17- and IFN-γ-producing γδ T cells

**DOI:** 10.1101/802728

**Authors:** Oliver Dienz, Victoria L. DeVault, Shawn C. Musial, Somen K. Mistri, Linda Mei, Aleksandr Baraev, Julie A. Dragon, Dimitry Krementsov, Andre Veillette, Jonathan E. Boyson

## Abstract

During thymic development, γδ T cells commit to either an IFN-γ- or an IL-17-producing phenotype through mechanisms that remain unclear. Here, we investigated whether the SLAM/SAP signaling pathway played a role in the functional programming of thymic γδ T cells. Characterization of SLAM family receptor expression revealed that thymic γδ T cell subsets were each marked by distinct co-expression profiles of SLAMF1, SLAMF4, and SLAMF6. In the thymus, immature CD24^hi^ Vγ1 and Vγ4 γδ T cells were largely contained within a SLAMF1^+^SLAMF6^+^ double positive (DP) population, while mature CD24^low^ subsets were either SLAMF1^+^ or SLAMF6^+^ single positive (SP) cells. In the periphery, SLAMF1 and SLAMF6 expression on Vγ1, Vγ4, and Vγ6 T cells distinguished IL-17- and IFN-γ-producing subsets, respectively. Disruption of SLAM family receptor signaling through deletion of SAP resulted in impaired thymic γδ T cell maturation at the CD24^hi^SLAMF1^+^SLAMF6^+^ DP stage that was associated with a decreased frequency of CD44^+^RORγt^+^ γδ T cells. These defects were in turn associated with impaired γδ T cell IL-17 and IFN-γ production in both the thymus as well as in peripheral tissues. The role for SAP was subset-specific, as Vγ1, Vγ4, Vγ5, but not Vγ6 subsets were SAP-dependent. Together, these data suggest that the SLAM/SAP signaling pathway regulates a critical checkpoint in the functional programming of IL-17 and IFN-γ-producing γδ T cell subsets during thymic development.

## Introduction

Innate-like γδ T cells are unusual T cells that are highly enriched in skin and mucosal tissues where they constitute a prominent source of cytokines and chemokines (1). In mice, most γδ T cells produce either IL-17 or IFN-γ, and IL-17 production is primarily restricted to the CD27^−^CCR6^+^ Vγ4 and Vγ6 subsets (Heilig and Tonegawa nomenclature (2)), whereas IFN-γ production is observed in CD27^+^CCR6^−^ Vγ1, Vγ4, Vγ5, and Vγ7 subsets (3–6). Accumulating data suggest that the relative balance of these IL-17 or IFN-γ-producing subsets can have significant consequences on disease outcome. IFN-γ-producing γδ T cells, for example, contribute to anti-tumor immunity (7) and inflammation in a model of cerebral malaria (3, 8). IL-17-producing γδ T cells, in contrast, have been associated with protection against bacterial and fungal infections (9–11), and promotion of tumor metastasis (12, 13) and immunopathologies in settings of autoimmunity (14–19).

Like their innate-like T cell counterparts, iNKT cells and MAIT cells, the functional programming of innate-like γδ T cells occurs during thymic development. Unlike iNKT and MAIT cells, however, a significant portion of γδ T cell developmental programming is restricted to a discrete developmental window (20), as it has been demonstrated that most natural IL-17-producing γδ T cells (γδT17) are generated only during the embryonic/early post-natal stage (21). Although the mechanisms that regulate the differentiation of γδ T cells into IFN-γ or IL-17-producing subsets remain unclear, a significant body of data supports a model in which strong or weak TCR signals skew the developing γδ T cell toward IFN-γ (strong signal) or IL-17 (weak signal) production (22–26). In contrast, there is also evidence supporting a TCR-independent model in which the development of some γδT17 subsets is dependent on pre-existing hardwired transcriptional programs (27–29). Exactly how these seemingly independent mechanisms converge to regulate the developmental programming of innate-like γδ T cell subsets remains an open question.

Slam receptors comprise a family of nine cell surface receptors, SLAMF1 (SLAM; CD150), SLAMF2 (CD48), SLAMF3 (Ly9; CD229), SLAMF4 (2B4; CD244), SLAMF5 (CD84), SLAMF6 (Ly108; CD352, SLAMF7 (CRACC; CD319), SLAMF8 (BLAME), and SLAMF9 (SF2001; CD84H), that are expressed primarily on hematopoietic cells and which play diverse roles in immune development and function (30). Most Slam receptors interact in a homophilic manner and therefore serve as self-ligands, the exception to this rule being the heterophilic interaction between SLAMF4 and SLAMF2. SLAM receptor signals can be both activating or inhibitory, depending on balanced recruitment of the cytosolic signaling adapter proteins SAP (31) and EAT-2 (32), and inhibitory SHP-1 or SHIP phosphatases to immunoreceptor tyrosine switch motifs in the SLAM family receptor cytoplasmic domain.

Inactivation of the gene encoding SAP (*Sh2d1a*) disrupts the signaling pathway of the majority of SLAM family receptors and results in the development of the primary immunodeficiency X-linked lymphoproliferative disease (XLP) which is characterized by broad immunological defects, among which is a failure in iNKT cell development (33, 34). Interestingly, the only other lymphocyte subset in mice whose development is dependent on SAP expression is the Vγ1Vδ6.3 (γδNKT) T cell subset (35), indicating that the requirement for SLAM/SAP signaling during development is shared by NKT cells and at least one γδ T cell subset. Yet, very little is known regarding SLAM family receptor expression on γδ T cells in either the periphery or the thymus. In this study, therefore, we explored whether the SLAM/SAP signaling pathway played a role in the developmental programming of thymic γδ T cells that determines their effector function.

## Materials and Methods

### Mice

C57BL/6J, BALB/cJ, and B6.*Sh2d1a^−/−^* (SAP^−/−^) mice were purchased from The Jackson Laboratory (Bar Harbor, ME). Mice were housed and bred in a specific pathogen-free barrier room at the AAALAC-approved animal facility of the University of Vermont. Age and sex matched mice between 8 and 12 weeks of age were used for experiments involving adult naïve mice as indicated in the figure legends. To analyze γδ T cells in embryonic thymus, timed-pregnant matings were used. The presence of a vaginal plug indicated day 0.5 of pregnancy. No attempt of sex matching was done for fetuses and newborn mice. All procedures involving animals were approved by the University of Vermont Institutional Animal Care and Use Committee.

### Cell preparation and stimulation

Single cell suspensions of splenocytes, lymph nodes, and thymocytes were obtained by gently pressing tissue through nylon mesh. RBCs were lysed using Gey’s solution. Lung cell preparations were conducted as described (36). Lungs were dissected away from the trachea and finely minced in enzymatic digestion buffer (Dulbecco’s modified Eagle’s medium (DMEM), 1 mg/ml collagenase type IV (Invitrogen), 0.2 mg/ml DNase (Sigma)). Lung tissue was resuspended in additional enzymatic digestion buffer and shaken at 200 rpm at 37°C for 20 min. The samples were triturated with a 16-gauge needle and allowed to digest for an additional 20 min, after which they were triturated one last time. After RBC lysis using Gey’s solution, the digested tissue was passed through a 70-μm filter and washed in PBS/2% FCS.

Single cell suspensions from skin were prepared as described (37). Briefly, depilated skin was digested in Trypsin/GNK buffer (0.3% bovine pancreatic trypsin (Type XI) (Sigma), 5 mM KCl, 150 mM NaCl, and 0.1% glucose, pH 7.6) for 2 h at 37 °C., after which the epidermis was separated from the dermis. Epidermis was then placed in Trypsin/GNK buffer containing 0.5% DNase I, and incubated for 15 min at 37 °C. After incubation, samples were passed through 70 μm mesh, washed with DMEM + 10% FBS, and resuspended in RPMI 1640 supplemented with 2 mM l-glutamine, 50 μM 2-mercaptoethanol, 50 U/ml penicillin, 50 ug/ml streptomycin + 10 U/ml IL-2 and incubated overnight at 37 °C in a 5% CO_2_ incubator prior to staining.

Liver intrahepatic leukocytes (IHLs) were isolated as previously described (38). Briefly, livers were perfused through the portal vein with PBS, subsequently removed, minced, and gently pressed through nylon mesh. The resulting cell suspension was washed twice and then resuspended in isotonic 33.8% Percoll (GE Healthcare). After centrifugation, the IHL cell pellet was resuspended, washed with PBS, and red blood cells were lysed using Gey’s solution. Single cell suspensions were resuspended in flow cytometry staining buffer.

Intestinal Intraepithelial Lymphocytes (IEL) were isolated as previously described (39). Briefly, mouse small intestines were flushed with cold CMF buffer (1 mM Hepes, 2.5 mM NaHCO_3_, 2% FBS in Ca^2+^/Mg^2+^-free HBSS pH 7.2). Surrounding fat tissue was removed, intestines opened with a longitudinal incision, and excess mucous removed by gentle scraping. Tissue was cut into small pieces, washed 3 times with CMF, followed by shaking in CMF+10% FBS/1 mM Dithioerythritol (DTE) for 20 minutes at 37 °C. This step was repeated and the cells in the supernatants of both extractions were combined. Cells were incubated for 20 minutes at 4 °C, before transferring cell supernatants into a fresh tube. Cells were centrifuged, washed once with RPMI, before resuspending in 44% Percoll solution. Cells were overlaid on a 67% Percoll solution, spun for 20 minutes at room temperature. Cells at the interface were collected, washed twice with RPMI, prior to counting and further analysis.

For all tissues, cell numbers were determined on a MACSquant VYB (Miltenyi) using PI (1 ug/ml in PBS) for dead cell exclusion. Cell stimulations were conducted in Bruff’s medium (Click’s medium (Sigma) supplemented with 2 mM l-glutamine, 50 μM 2-mercaptoethanol, 50 μg/ml gentamycin, 50 U/ml penicillin, 50 ug/ml streptomycin, and 5% FBS) with 25 ng/ml PMA and 500 ng/ml ionomycin for 4 hours at 37 °C and 5% CO_2_. After 2 hours, 2 μM monensin was added to the culture.

### Flow Cytometry

Cells were stained with Live Dead Fixable Blue staining reagent (ThermoFisher, Grand Island, NY) at 4°C in PBS for 30 min., after which they were washed and resuspended in PBS/2% FCS containing 0.1% sodium azide and Fc-Block (Biolegend) to block non-specific antibody binding prior to the addition of antibodies. Brilliant Violet Staining buffer (BD Biosciences) was used in experiments containing more than one Brilliant Violet or Brilliant Ultraviolet fluorophore conjugated antibody. To identify the Vγ6 subset, the staining procedure described in Roark et al. was followed (40).

Cells were washed twice with PBS/2% FCS/0.1% sodium azide solution and fixed in PBS/1% PFA buffer. For intracellular cytokine staining, cells were permeabilized with PBS/1% FCS/0.1% sodium azide/0.1% saponin after surface staining and fixation. 25 μg/ml total rat IgG were added to reduce unspecific binding prior to cytokine staining with fluorescent-labeled antibodies. Nuclear transcription factors were analyzed using the Foxp3 transcription factor fixation/permeabilization reagent (ThermoFisher). Briefly, after staining for surface markers cells were resuspended in the fixation/permeabilization reagent and incubated overnight at 4 °C. Cells were washed, incubated with 25 μg/ml total rat IgG prior to adding antibodies against transcription factors. After staining at 4 ^ο^C, cells were washed and resuspended in PBS/2% FCS/0.1% sodium azide. Flow cytometry data were collected on a LSRII flow cytometer (BD Bioscience) and analyzed with FlowJo software (Treestar).

Antibodies used in these experiments were: CD11b (M1/70), CD19 (1D3), CD24 (M1/69), CD27 (LG.3A10), CD44 (IM7), CD45 (30-F11), CD48 (HM48-1), CD84 (mCD84.7), CD150 (TC15-12F12.2), CD229 (Ly9ab3), CD352 (330-AJ), IFN-g (XMG1.2), IL-17A (TC11-18H10.1), TCRβ (H57-597), TCRδ (GL3), Vγ1 (2.11), Vγ5 (536) from Biolegend; CD196 (140706), CD244 (2B4), RORγt (Q31-378) from BD Biosciences; CD244.1 (C9.1), CD352 (13G3-19D), Eomes (Dan11mag), Vγ4 (UC3-10A6) from eBiosciences.

### t-SNE analysis

Dimensionality reduction of flow cytometry data was conducted using t-stochastic neighbor embedding (t-SNE) analysis in FlowJo. First, traditional gating was used to identify γδ T cells, after which the γδ T cell node data from 5 individual mice was concatenated and exported for further analysis. t-SNE analysis was conducted on the SLAMF1, SLAMF4, SLAMF6, Vγ1, Vγ4, Vγ6, CD44, and CD27 parameters using the following conditions: 1000 iterations, perplexity = 40, learning rate = 436.

### Transcriptional profiling

Leukocytes from lung cell preparations were isolated using anti-CD45 Microbeads and LS columns according to the manufacturer’s instruction (Miltenyi). Cells were stained with Fixable Viability Dye eFluor780 (eBioscience), washed, and surface stained with Abs against SLAMF1, SLAMF6, NK1.1, TCRδ, TCR-Vγ1, TCR-Vγ4, TCRβ, CD11b, CD19, and CD45 in sterile PBS/2% FBS. Cells were sorted on a FACS Aria III (BD Bioscience) into culture medium, pelleted, washed, and snap frozen until later use. RNA was isolated using a RNeasy Micro kit (Qiagen). RNA quality was assessed using an Agilent Bioanalyzer 2100, and RNA concentration was calculated using a Qubit fluorometer (ThermoFisher Scientific). Amplified cDNA was prepared using the Ovation Pico WTA System V2 (Nugen), and biotinylated using the Encore biotin module (Nugen). Hybridization with GeneChip Mouse Gene 2.0 ST arrays (Affymetrix) was conducted overnight at 45 °C. at 60 rpm. CEL files were analyzed using Transcriptome Analysis Console (ThermoFisher) software. Genes that demonstrated a greater than or less than 2-fold difference in expression and a false-discovery rate of <0.2 were identified. This resulted in 316 SLAMF1-enriched genes and 229 SLAMF6-enriched genes. These gene lists were filtered by removing unannotated transcripts, predicted transcripts, and by removing all but 1 of the 28 multiple transcript isoforms of *Syne1*, yielding 244 SLAMF1-enriched and 181 SLAMF6-enriched transcripts. All differentially expressed genes in this list exhibited a p-value < 0.004. The raw data files for this analysis were deposited in NCBI GEO (GSE128544).

Data were analyzed for gene ontological and pathway enrichment using NIH DAVID analysis and IPA (filtered to include only those with a p-value and FDR <0.05). Heatmaps were generated using prettyheatmap with row-scaling. Upstream regulator prediction using IPA analysis was conducted on the 425 differentially expressed genes based on fold-change (SLAMF1^+^SLAMF6^−^/SLAMF1^−^SLAMF6^+^). The result was filtered to include only those predicted regulators with p-value of overlap < 1 × 10^−5^, ranked according to activation z-score, after

### Statistical analysis

Statistical analysis was performed using Prism (GraphPad Software, San Diego, CA). Unpaired Student t-tests, one-way ANOVA, or two-way ANOVA was used where appropriate. For ANOVA posttest comparisons, the Dunnett, Tukey, or Sidak multiple comparison tests were used where appropriate. In all cases, tests were considered significant when p < 0.05. Unless otherwise indicated, error bars represent standard deviation.

## Results

### Heterogeneous SLAM receptor expression profiles on thymic γδ T cell subsets

γδ T cell effector functions are largely programmed during thymic development, and IL-17 functional programming is thought to occur primarily during embryonic and neonatal thymic development (21). Therefore, we examined the co-expression of SLAMF1 through SLAMF6 receptors on γδ T cell subsets from C57BL/6 embryonic day 17 (E.17) thymus. This analysis revealed that SLAMF5 was not expressed on γδ T cells and that the expression of SLAMF2 and SLAMF3 was relatively uniform among γδ T cell subsets (**Fig. 1A**). In contrast, our analysis revealed that the other three receptors SLAMF1, SLAMF4, and SLAMF6 exhibited a surprising degree of heterogeneity in their co-expression (**Fig. 1B**). For example, a significant fraction of E.17 thymic γδ T cells were SLAMF1^−^SLAM4^+^SLAMF6^−^ and this population consisted primarily of Vγ5 γδ T cells (**Fig. 1C**). Mature Vγ5 in the skin exhibit the same SLAMF1^−^ SLAMF4^+^SLAMF6^−^ phenotype, suggesting that the Vγ5 SLAM receptor expression profile is established during embryonic thymic development (**Fig. 1D**) (41). Likewise, we observed that Vγ6 γδ T cells were primarily SLAMF1^+^SLAMF4^−^SLAMF6^−^ in the E.17 thymus, similar to the SLAM receptor expression profile that they exhibit in the adult lung (**Figs. 1B and 3A**).

**Fig. 1.**
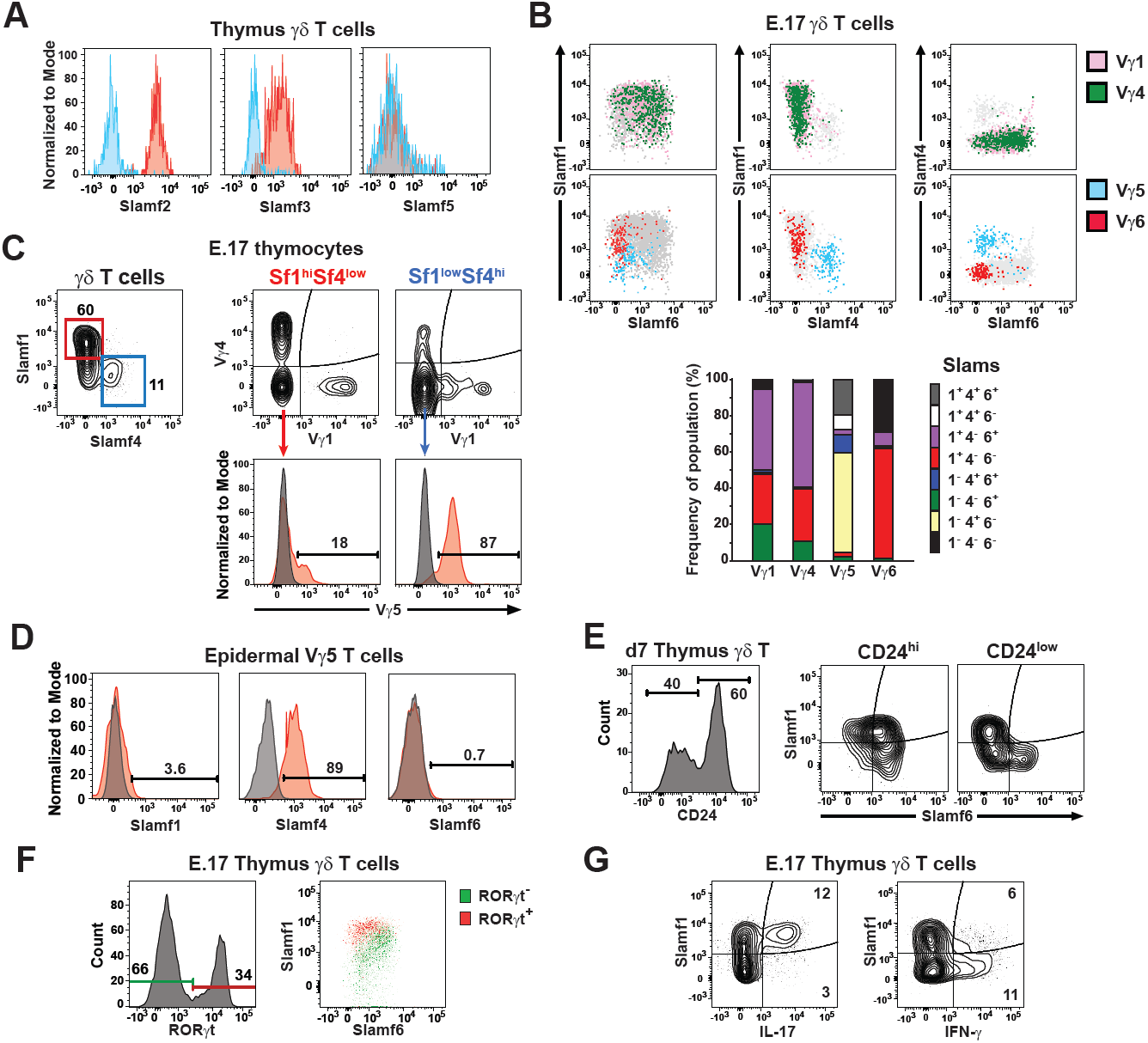
Slam receptor expression profiles on different γδ T cell subsets are established during thymic development. A) Expression pattern of SLAMF2, SLAMF3, and SLAMF5 on thymic E.17 γδ T cells. Red histograms indicate stained cells, while blue histograms indicate negative controls. Data are representative of 2 separate experiments, 4 mice per group, sex was not determined. B) SLAMF1, SLAMF4, and SLAMF6 expression on B6 E.17 γδ T cells subsets. The position of Vγ1^+^ (pink) and Vγ4^+^ (green) γδ T cells on plots of SLAMF1, SLAMF4, and SLAMF6 is shown above. The position of Vγ5^+^ (blue) and Vγ6^+^ (red) γδ T cells is shown below. Total γδ T cells are indicated in grey. Below, cumulative frequencies of Slam receptor expression profiles on the thymic γδ T cell subsets are shown as bar graph. Data are representative of 2 independent experiments, 6 mice each, sex was not determined. C) SLAMF1 and SLAMF4 expression on E.17 thymic Vγ1, Vγ4, and Vγ5 subsets. Representative contour plots of SLAMF1 and SLAMF4 staining on thymic γδ T cells is shown at left. SLAMF1^hi^SLAMF4^lo^ and SLAMF1^lo^SLAMF4^hi^ γδ T cells are gated in red and blue, respectively. Vγ1, Vγ4, and Vγ5 staining on SLAMF1^hi^SLAMF4^lo^ and SLAMF1^lo^SLAMF4^hi^ subsets is shown at right. Data is representative of 2 independent experiments, 6-7 mice each, sex was not determined. D) SLAMF1, SLAMF4, and SLAMF6 staining on Vγ5^+^ dendritic epidermal T cells isolated from adult male B6 mice. Data are representative of 2 independent experiments. E) SLAMF1 and SLAMF6 expression on CD24^hi^ and CD24^low^ neonatal (day 7) thymus γδ T cells. Numbers indicate the frequency of gated cells. Data represent 2 independent experiments, 5 and 7 mice each, sex was not determined. F) SLAMF1 and SLAMF6 expression on RORγt^+^ and RORγt^−^ E.17 thymic γδ T cells. The position of RORγt^+^ (red) and RORγt^−^ (green) cells are shown on a dot plot of SLAMF1 and SLAMF6 expression. Data are representative of 3 independent experiments with 3-8 mice each experiment, sex was not determined. G) SLAMF1 expression on IL-17 and IFN-γ-producing E.17 thymic γδ T cells. Representative contour plots are shown. Data are representative of 2 independent experiments with 4 and 6 mice each experiment, sex was not determined.

**Fig. 2.**
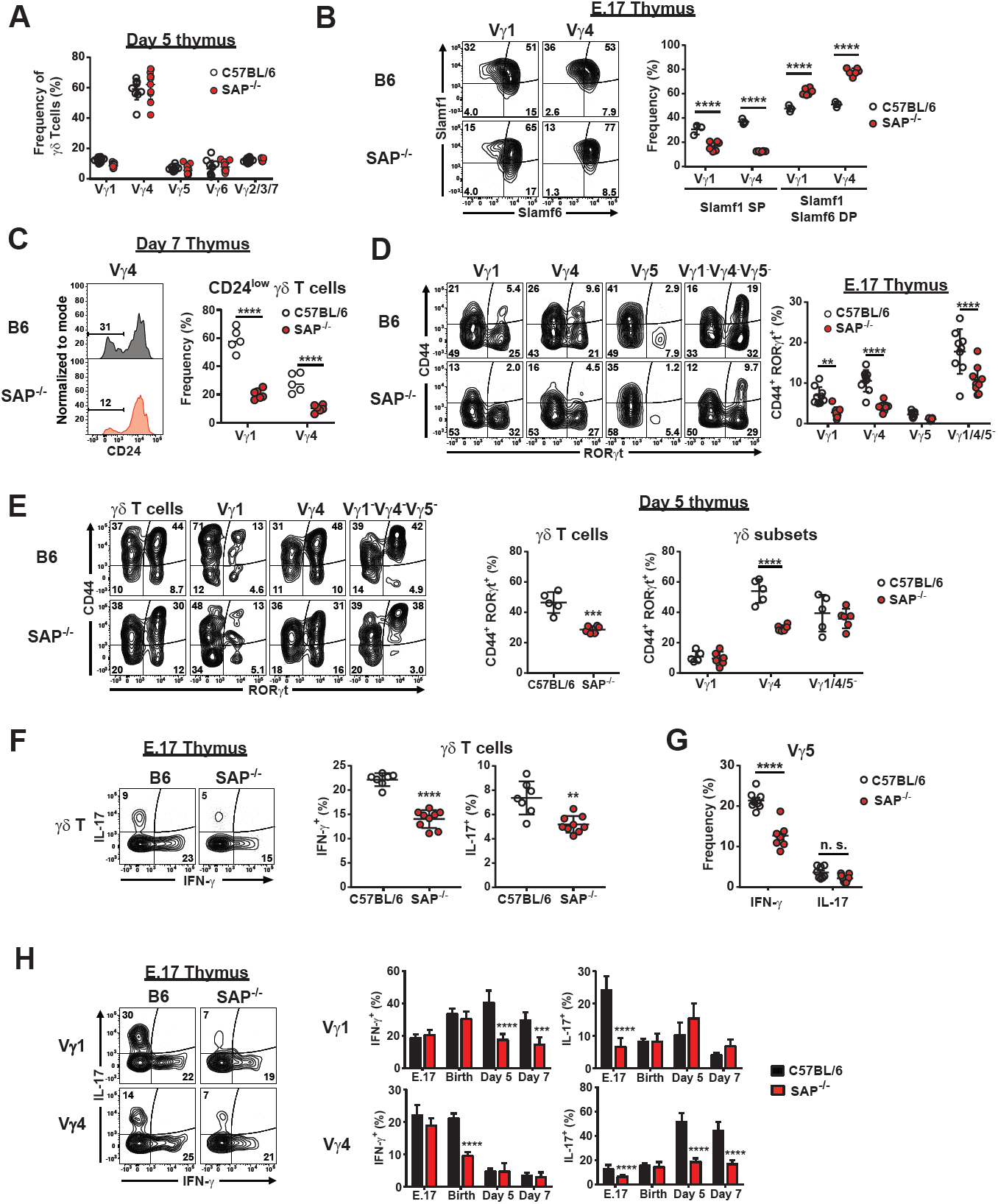
Impaired γδ T cell functional programming in SAP-deficient mice. A) Frequencies of γδ T cell subsets from day 7 C57BL/6J and B6.SAP^−/−^ thymus. The combined data from 2 independent experiments is shown, sex was not determined. B) SAP-dependent alteration in SLAMF1 SP and SLAMF1SLAMF6 DP γδ T cells. Representative contour plots depicting SLAMF1 and SLAMF6 expression on E.17 and day 6 thymic Vγ4 T cells is shown at left. The frequencies of SLAMF1 and SLAMF6-expressing subsets is shown at right, ****p < 0.0001, as determined by two-way ANOVA followed by Sidak’s multiple comparisons test. Sex of the mice was not determined. C) Decreased CD24^low^ Vγ1 and Vγ4 T cells in SAP-deficient mice. Representative histograms depicting CD24 expression on day 7 thymic Vγ4 cells are shown at left. Data represent 2 independent experiments, 5-7 mice per group, sex was not determined. ****p < 0.0001, as determined by two-way ANOVA followed by Sidak’s multiple comparisons test. D) Significant decrease in CD44^+^RORγt^+^ thymic γδ T cells in E.17 B6.SAP^−/−^ mice. Thymocytes from E.17 B6 and B6.SAP^−/−^ mice were stained for surface markers and nuclear staining was used to assess RORγt expression. Representative contour plots depicting CD44 and RORγt expression are shown at left. Cumulative frequencies of CD44^+^RORγt^+^ E.17 thymic γδ T cells is shown at right, **p < 0.01, ****p < 0.0001, as determined by two-way ANOVA followed by Sidak’s multiple comparisons test. Data are representative of 2 independent experiments, 7-9 mice per group, sex was not determined. E) SAP-dependent decrease in CD44^+^RORγt^+^ γδ T cells in day 5 neonate thymus. Representative contour plots are shown at left. Cumulative frequencies of CD44^+^RORγt^+^ are shown at right, ***p < 0.001, ****p < 0.0001, as determined by two-way ANOVA followed by Sidak’s multiple comparisons test. Data are representative of 2 independent experiments, 5-6 mice per group, sex was not determined. F) Intracellular IFN-γ and IL-17 production in E.17 thymic γδ T cells. Representative contour plots are shown at left. Numbers indicate the percentage of cells. Cumulative data are shown at right, **p < 0.01, ****p < 0.0001 as determined by an unpaired two-tailed Student’s t-test. Sex of the mice was not determined. G) Decreased E.17 Vγ5 IFN-γ production in SAP-deficient mice. ****p < 0.0001, as determined by Student’s t-test, sex was not determined. H) IFN-γ and IL-17 production in thymic γδ T cell subsets from B6 and B6.SAP^−/−^ mice at different developmental stages as indicated. Cumulative data represent the combined results of 4 independent experiments, 5 – 9 mice per group, sex was not determined. ***p < 0.001, ****p < 0.0001, as determined by two-way ANOVA followed by Sidak’s multiple comparisons test.

**Figure 3.**
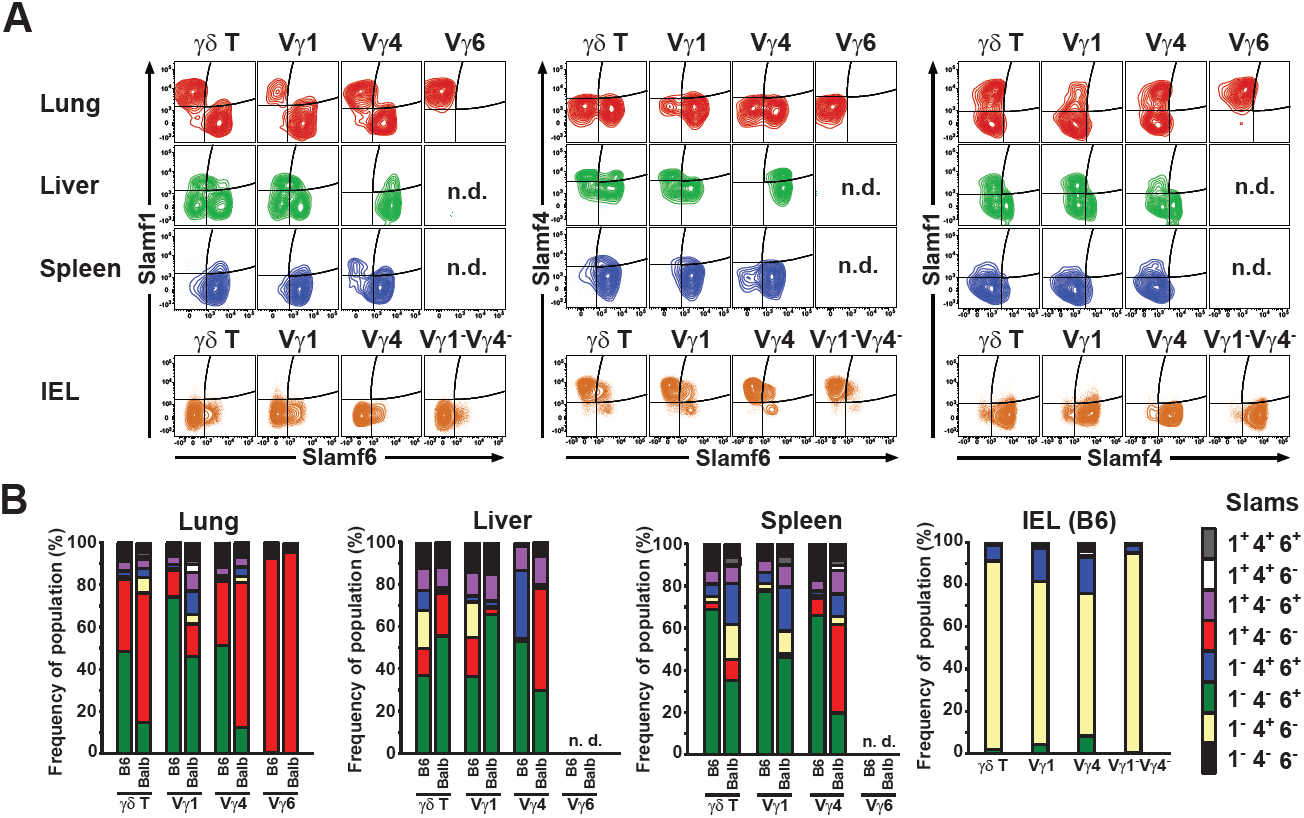
Tissue-dependent heterogeneity in γδ T cell SLAM family receptor expression profiles. A) Representative contour plots of SLAMF1, SLAMF4, and SLAMF6 expression on lung, liver, spleen, and IEL γδ T cells. Data represent concatenated files from 4 - 5 female B6 mice 11 weeks of age. n.d., not detected. B) Cumulative frequencies of Slam receptor expression profiles on γδ T cell subsets from B6 and BALB/c mice. Data represent groups of 4 - 5 B6 mice (as in A) and 4 female BALB/cJ mice, 12 weeks of age. IEL γδ T cells from BALB/cJ mice were not analyzed.

In contrast to the Vγ5 and Vγ6 populations, E.17 Vγ1 and Vγ4 exhibited broad expression of SLAMF1 and SLAMF6 and they were uniformly SLAMF4^−^ (**Fig. 1B**). We identified both SLAMF1 and SLAMF6 single positive (SP) populations that were present in peripheral tissues, as well as a SLAMF1SLAMF6 double positive (DP) population that was relatively rare in the periphery (**Fig. 1B and Fig. 3A**). Interestingly, further investigation revealed that the SLAMF1SLAMF6 DP population was largely contained within the CD24^hi^ γδ T cell population, while the SLAMF1 and SLAMF6 SP populations, were primarily CD24^low^. These observations suggested that immature CD24^hi^ γδ T cells exhibited a SLAMF1SLAMF6 DP phenotype that resolved into either CD24^low^ SLAMF1 or SLAMF6 SP phenotypes or a SLAMF1SLAMF6 DN upon maturation (**Fig. 1E**).

Since it was previously reported that IL-17-producing γδ T cells preferentially express SLAMF1 (42), we asked whether the association of SLAMF1 receptor expression with IL-17 was established during thymic development. Interestingly, we found that RORγt^+^ E.17 γδ T cells were largely found in a SLAMF1^hi^SLAMF6^low^ γδ T cell subset (**Fig. 1F**) and that majority of IL-17-producing γδ T cells were SLAMF1^+^, while the majority of IFN-γ-producing γδ T cells were SLAMF1^−^ (**Fig. 1G**). Taken together, these data suggested that the co-expression of SLAMF1, SLAMF4, and SLAMF6 distinguished 3 groups of γδ T cell subsets at E.17: Vγ1/Vγ4, Vγ5, and Vγ6. Moreover, the data suggested that immature thymic Vγ1 and Vγ4 γδ T cells initially possess a SLAMF1SLAMF6 DP profile, and then develop into either SLAMF1 or SLAMF6 single positive expression profiles concomitant with maturation and acquisition of function.

### Impaired γδ T development in SAP-deficient mice

SLAMF1, SLAMF4, and SLAMF6 transduce signals by utilizing the signaling adapter proteins SAP and EAT-2, which binds to immunoreceptor tyrosine switch motifs (ITSMs) located in the cytoplasmic tails of SLAM family receptors (30). *Sh2d1a* (encoding SAP) is the most widely expressed of these adapter proteins in γδ T cells and SAP-deficient mice have previously been reported to be deficient in the Vγ1Vδ6.3 (γδNKT) subset (43). To determine whether the SLAM/SAP signaling pathway played a role in the functional programming that γδ T cells undergo during thymic development, we examined γδ T cell development in SAP-deficient mice during embryonic and neonatal stages of development.

A comparison of thymic γδ T cells between B6 and B6.SAP-deficient mice revealed no significant change in the distribution of Vγ1, Vγ4, Vγ5, or Vγ6 γδ T cell subsets (**Fig. 2A**), although we did confirm the critical role of SAP in the development of the Vγ1Vδ6.3 subset (**Suppl. Fig. 1**). An examination, however, of the SLAM receptor expression profiles on E.17 γδ T cells revealed a significant decrease in the frequency of SLAMF1 SP Vγ1 and Vγ4 T cells (**Fig. 2B**). The reduction in the SLAMF1 SP subset coincided with a significant increase in the frequency of SLAMF1SLAMF6 DP subsets in B6.SAP^−/−^ mice both at E.17 (**Fig. 2B**) and at day 6 (**Suppl. Fig. 1**). In accordance with our previous observation that SLAMF1SLAMF6 DP and SLAMF1 and SLAMF6 SP γδ T cells were CD24^hi^ and CD24^low^, respectively (**Fig. 1D**), we observed a significant decrease in the frequency of CD24^low^ thymic Vγ1 and Vγ4 γδ T cells in SAP-deficient mice (**Fig. 2C**). No effect of SAP deficiency in CD24 expression on Vγ6 subsets was observed (not shown). Together, these data suggested that SAP plays a role in the maturation of Vγ1 and Vγ4 γδ T cells during thymic development.

Next, we assessed whether SAP-deficient thymic γδ T cells exhibited any functional defects. Examination of the E.17 CD44^+^RORγt^+^ γδ T cell population revealed a significant decrease in the frequency of CD44^+^RORγt^+^ γδ T cells in SAP-deficient mice (**Fig. 2D**). Significant decreases were observed in the Vγ1, Vγ4, and Vγ1^−^4^−^5^−^ subsets, but not Vγ5, very few of which were CD44^+^RORγt^+^ (**Fig. 2D**). Examination of neonatal day 5 thymocytes revealed that the SAP-dependent decrease in CD44^+^RORγt^+^ γδ T cells was not restricted to embryonic thymic development, as there was a significant decrease in the frequency of CD44^+^RORγt^+^ Vγ4 T cells, which comprise the majority of thymic γδ T cells at this stage of development (**Fig. 2E**). However, the SAP-dependent decrease in CD44^+^RORγt^+^ Vγ1 and Vγ1^−^ 4^−^5^−^ (primarily Vγ6 at this stage of development) populations was not observed in the neonatal thymus (**Fig. 2E**).

Finally, we investigated whether SAP played a role in the acquisition of γδ T cell function during thymic development. A comparison of E.17 γδ T cell cytokine production between B6 and B6.SAP^−/−^ mice revealed a significant decrease in both IL-17 and IFN-γ production in SAP-deficient mice (**Fig. 2F**). Subset analysis revealed that the reduction in IFN-γ production was due primarily to an effect on Vγ5 γδ T cells (**Fig. 2G**), while the reduction in IL-17 production was instead observed in both Vγ1 and Vγ4 γδ T subsets (**Fig. 2H**). No significant decrease in the IL-17-producing Vγ1^−^Vγ4^−^Vγ5^−^ population was observed at this time, suggesting that development of the Vγ6 population was SAP-independent (data not shown).

Analysis of later stages of development revealed that the effect of SAP deficiency was dependent on both the subset and developmental stage. A significant effect of SAP deficiency was observed on thymic Vγ4 IL-17 production at all stages examined except for day 0. Conversely, we found that Vγ4 IFN-γ production was significantly decreased in SAP-deficient mice on day 0, but not any of the other timepoints examined. Similarly, we noted decreased thymic Vγ1 IL-17 in B6.SAP^−/−^ mice only at E.17, and decreased thymic Vγ1 IFN-γ only on days 5 and 7 (**Fig. 2H**). Together, these data suggested that the SLAM/SAP signaling pathway played a significant role in the developmental programming of thymic Vγ1, Vγ4, and Vγ5 IL-17 and IFN-γ production during embryonic and neonatal thymic development. In addition, the data suggested that impairments in the developmental programming of γδ T cells was linked to a SAP-dependent role in γδ T cell maturation.

### Tissue-specific heterogeneity in SLAM receptor expression profiles on γδ T cell subsets in the periphery

Next, we examined the extent to which SAP-dependent impairments in thymic γδ T cell development affected γδ T cell function in peripheral tissues. We first assessed the co-expression of SLAMF1 through SLAMF6 on γδ T cell subsets from different tissues. Similar to our results in the thymus, we found little evidence of SLAMF5 expression, and we found uniform expression SLAMF2 and SLAMF3 on γδ T cells and from the lung, spleen, and lymph node (**Suppl. Fig. 2**). In contrast, our analysis revealed a significant degree of heterogeneity in SLAMF1, SLAMF4, and SLAMF6 co-expression on γδ T cell subsets obtained from different tissues (**Fig. 3A**). In the lung, we found two distinct populations of Vγ1 and Vγ4 γδ T cells which were either SLAMF1 SP or SLAMF6 SP. In contrast, Vγ6 γδ T cells were uniformly SLAMF1 SP with no detectable SLAMF6 expression (**Fig. 3A**). A similar SLAM receptor expression pattern was observed in the spleen although the frequency of the SLAMF1 SP Vγ4 T cell population was markedly reduced compared with the lung and essentially absent in the Vγ1 subset (**Fig. 3A**). In contrast to lung and spleen, a significant proportion of SLAMF4-expressing γδ T cell subsets were detected in the liver, while gut IEL γδ T cells only expressed SLAMF4, and not SLAMF1 or SLAMF6 (**Fig. 3A**).

Heterogeneity in SLAM family receptor co-expression was also observed in BALB/cJ γδ T cells, which possess a different *Slam* haplotype (44) (**Fig. 3B**). We did note significant differences between the strains indicating that SLAM receptor co-expression profiles were influenced by genetic background. Notably, we observed a significantly greater proportion of SLAMF1^+^ SLAMF6^−^ lung and spleen Vγ4 T cells in BALB/c compared to their B6 counterparts (**Fig. 3B**). Together, these data indicated that the co-expression of SLAM family receptors on γδ T cells constituted unique SLAM receptor expression profiles that varied in both a tissue-dependent and Vγ subset-dependent manner. The data also indicated that the SLAM receptor expression profiles of γδ T cell subsets in the periphery largely reflect those established during thymic development, and suggested that they are relatively stable markers that result from a developmental program.

### SLAM receptor expression profiles distinguish γδ T cell functional subsets in the periphery

Given the association of SLAMF1 expression with γδ T cell IL-17 production in the thymus (**Fig. 1G**), we were interested in determining whether distinct SLAM receptor expression profiles marked functional subsets of γδ T cells in the periphery. We focused our attention on lung γδ T cells, since they exhibited a diversity of SLAM family receptor expression profiles. Analysis of the co-expression of SLAM family receptors with the surrogate functional markers, CD44 and CD27, on lung γδ T cells revealed that SLAMF1^+^SLAMF6^−^ γδ T cells were primarily CD44^+^CD27^−^ (**Fig. 4A**) consistent with an IL-17-producing phenotype (42). Interestingly, SLAMF1^−^SLAMF6^+^ γδ T cells were primarily CD44^−^CD27^+^ suggestive of an IFN-γ-producing phenotype, while SLAMF4-expressing γδ T cells were distributed in both CD44^+^CD27^−^ and CD44^−^CD27^+^ populations (**Fig. 4A**). Therefore, SLAMF1 and SLAMF6 expression in the periphery coincided with canonical markers of IL-17 and IFN-γ-producing γδ T cells, respectively.

**Figure 4.**
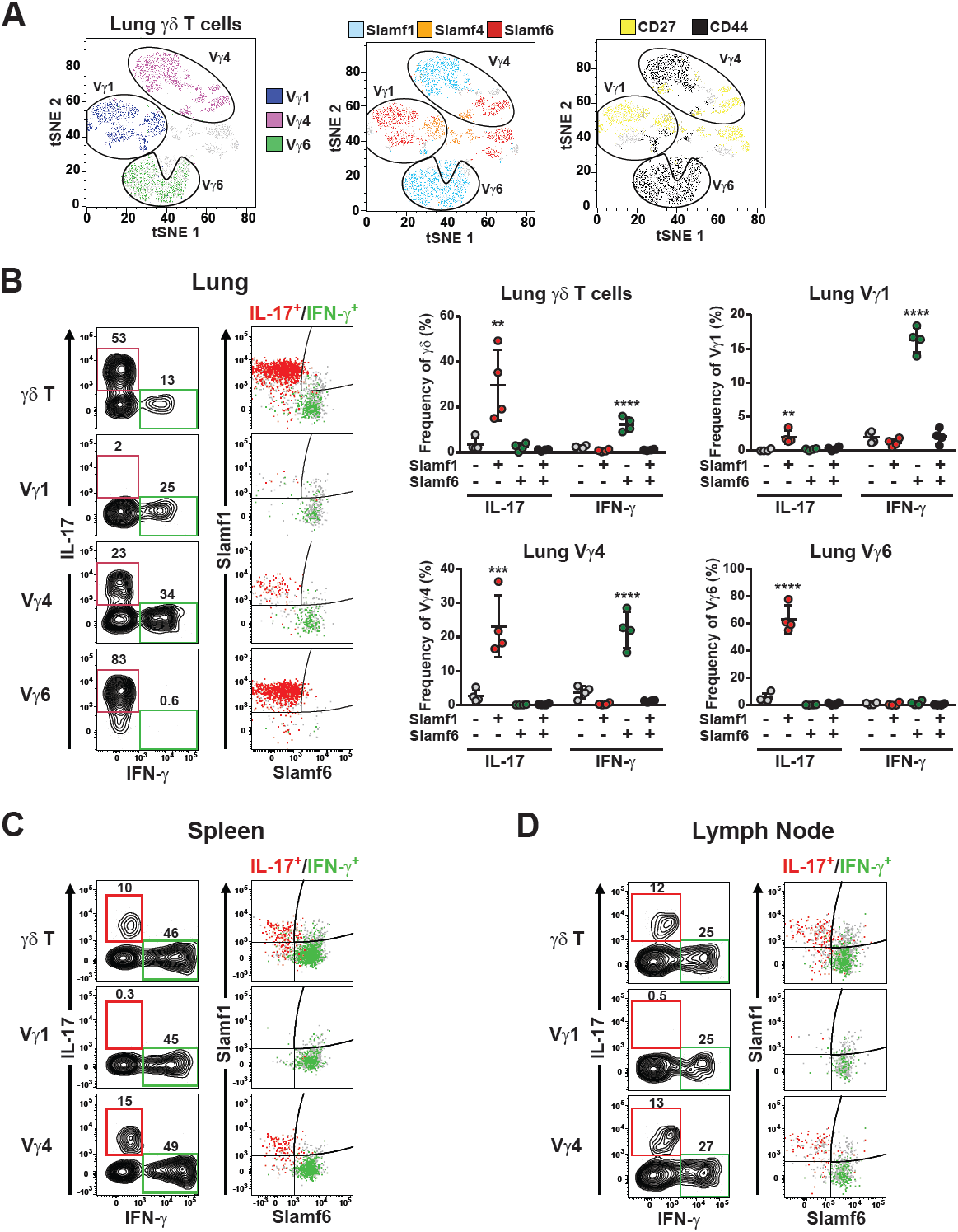
Slam receptor expression profiles distinguish distinct functional subsets of γδ T cells in the periphery. A) Slam family receptor co-expression segregates with surrogate markers of γδ T cell function. Flow cytometric data from concatenated from 5 individual female C57BL/6J mice 10 weeks of age was analyzed using t-SNE. Clusters representing Vγ1, Vγ4 and Vγ6 subsets are circled (Top), and SLAMF1, SLAMF4, and SLAMF6-expressing cells (bottom left), and CD44- and CD27-expressing cells (bottom right) are colored as indicated. B-D) SLAMF1 and SLAMF6 expression distinguish IL-17 and IFN-γ-producing γδ T cells in lung (B), spleen (C), and lymph node (D). Cell suspensions were stimulated with PMA/ionomycin for 4 h at 37 °C. in the presence of monensin, after which they were stained for surface markers and intracellularly stained for IFN-γ and IL-17. Representative contour plots of γδ T cell cytokine production after stimulation are shown at left. IL-17 and IFN-γ-producing cells are indicated by red and green gates, respectively. Dot plots depicting SLAMF1 and SLAMF6 expression are shown at right and the position of IL-17-gated and IFN-γ-gated cells are shown in red and green, respectively. All other cells are shown in grey. Cumulative data of lung γδ T cell cytokine production is shown at right. Data are representative of 3 independent experiments, 4 female C57BL/6J mice per group, 9.5 weeks of age,**p < 0.01, ***p < 0.001, ****p < 0.0001 as determined by one-way ANOVA followed by Tukey’s multiple comparison test.

To determine whether SLAM receptor expression profiles marked functional subsets of γδ T cells, we assessed lung γδ T cell cytokine production after stimulation with PMA/ionomycin. These data revealed that lung γδ T cell IFN-γ production was associated with SLAMF1^−^SLAMF6^+^ γδ T cells, while IL-17 production was associated with SLAMF1^+^SLAMF6^−^ γδ T cells (**Fig. 4B**). Similarly, stimulation of both spleen (**Fig. 4C**) and lymph node (**Fig. 4D**) γδ T cells revealed a strong association of SLAMF1 and SLAMF6 expression with γδ T cell IL-17 and IFN-γ production, respectively. Together, these data suggested that the mutually exclusive expression of SLAMF1 and SLAMF6 marked IL-17- and IFN-γ-producing Vγ1, Vγ4, and Vγ6 γδ T cell subsets in the periphery.

To explore this association further, we compared the *ex vivo* transcriptional profiles of sorted lung SLAMF1^+^SLAMF6^−^ and SLAMF1^−^SLAMF6^+^ lung Vγ4 from naïve C57BL/6J mice. This analysis revealed the differential expression of 244 genes in SLAMF1^+^SLAMF6^−^ and 181 genes in SLAMF1^−^SLAMF6^+^ lung Vγ4 (**Fig. 5A, Suppl. Table 1**). In accordance with our functional analysis, we found that SLAMF1^+^SLAMF6^−^ lung Vγ4 γδ T cells exhibited a transcriptional profile highly suggestive of IL-17 production. We observed differential expression of *Il23r, Il17a, Il17f, Il1r1, Blk, Rorc, Sox13, Maf, Ccr6, Ccr2, and Scart2* all of which have been previously associated with a γδT17 phenotype (27, 28, 45) (**Fig. 5A and Suppl. Table 1**). In contrast, the transcriptional profile of the SLAMF1^−^SLAMF6^+^ Vγ4 γδ T cells was suggestive of an IFN-γ-producing phenotype. *Ifng, Eomes, IL12rb, and Crtam* were preferentially expressed in SLAMF1^−^SLAMF6^+^ Vγ4 γδ T cells (**Fig. 5A and Suppl. Table 1**). These findings were consistent with a recent report demonstrating that EOMES expression marks IFN-γ-producing γδ T cells (46).

**Figure 5.**
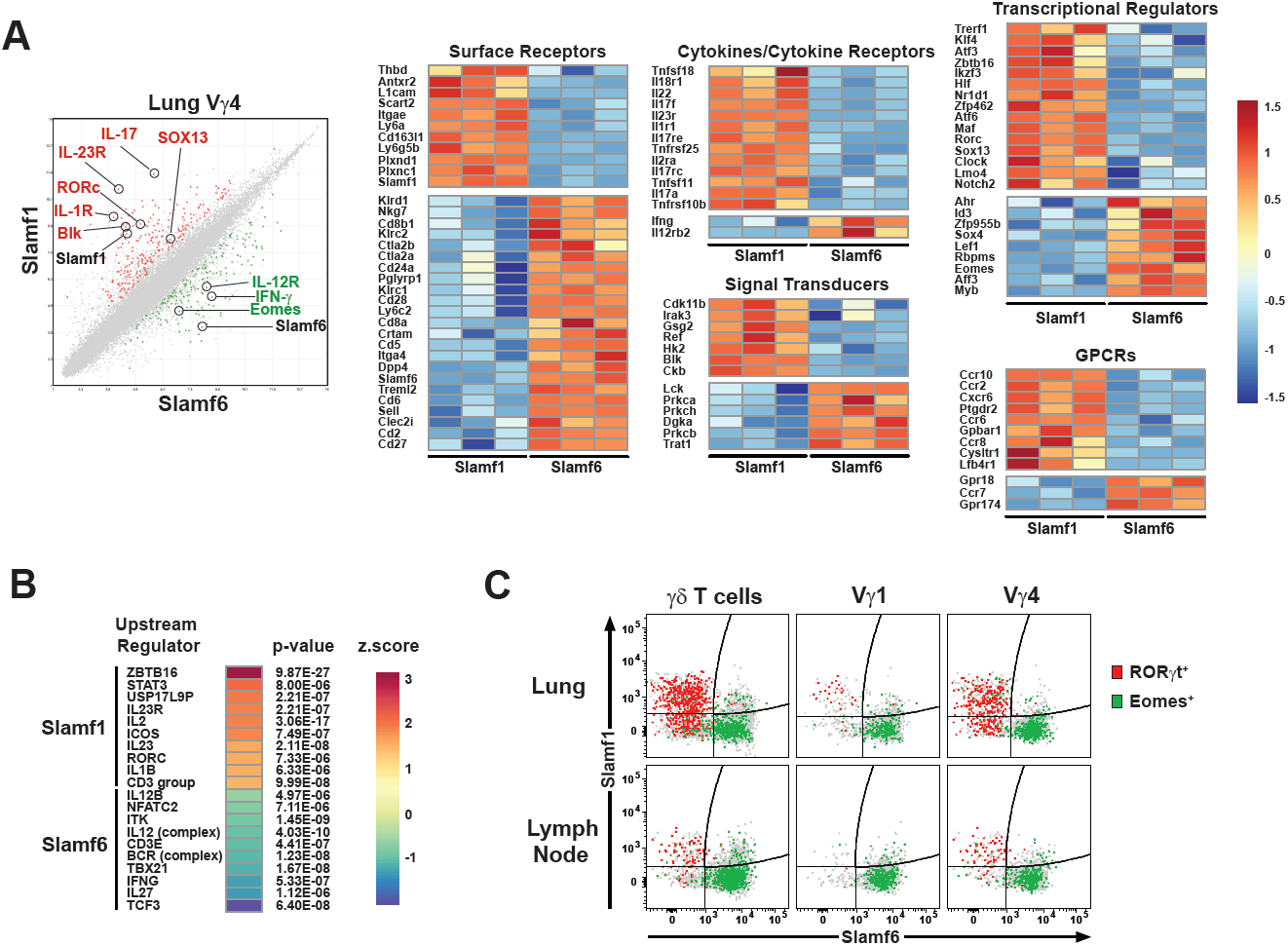
SLAMF1 and SLAMF6 expression distinguishes lung Vγ4 subsets with distinct transcriptional profiles. SLAMF1^+^SLAMF6^−^ and SLAMF1^−^SLAMF6^+^ lung Vγ4 cells were FACS-sorted from lung cell suspensions pooled from 12 female C57BL/6J mice each (n = 3 pools). RNA from sorted cells was used in a microarray to assess differential gene expression. A) Scatter plot of average sample signals (log_2_ transformed) observed in SLAMF1^+^SLAMF6^−^ and SLAMF1^−^SLAMF6^+^ cells is shown at left. Genes exhibiting a fold-change of greater or less than 2 and a false discovery rate < 0.2 are colored red (differentially expressed in SLAMF1^+^SLAMF6^−^) or green (differentially expressed in SLAMF1^−^SLAMF6^+^). Row-normalized heatmaps of differentially expressed genes are shown at right. Receptors, signal transducers, transcriptional regulators, cytokine/cytokine receptors, and GPCRs that exhibited a fold-change of greater or less than 2.5 are shown. B) Predicted upstream regulators of differentially expressed genes. Predicted regulators were ranked according to their activation z-score. Positive z-scores suggest activity in SLAMF1^+^SLAMF6^−^ and negative z-scores suggest activity in SLAMF1^−^SLAMF6^+^ subsets. C) SLAMF1 and SLAMF6 distinguish RORγt- and Eomes-expressing γδ T cell subsets. Dot plots depict SLAMF1 and SLAMF6 expression on lung and lymph node γδ T cell subsets. RORγt- and EOMES-expressing γδ T cells are highlighted in red and green, respectively. All other cells are shown in grey. Data are representative of two separate experiments, 4-5 male C57BL/6J mice, 13 weeks of age per group.

Pathway analysis (IPA) of differentially expressed genes revealed that transcripts enriched in SLAMF1^+^SLAMF6^−^ lung Vγ4 cells exhibited significant overlap with genes regulated by ZBTB16 (PLZF), IL23R, RORγT, and IL-1B. In contrast, transcripts enriched in SLAMF1^−^ SLAMF6^+^ lung Vγ4 exhibited significant overlap with genes regulated by IL-12, TBX21 (T-bet), IFN-γ, IL-27, and TCF3 (E2A) (**Fig. 5B and Suppl. Table 2**). These transcriptional profiles were confirmed using flow cytometry which revealed that RORγt expression was largely confined to the SLAMF1^+^SLAMF6^−^ γδ T cells in both lung and lymph node, while EOMES expression was found in the SLAMF1^−^SLAMF6^+^ γδ T cell subsets (**Fig. 5C**). Together, these data suggested that lung SLAMF1^+^SLAMF6^−^ and SLAMF1^−^SLAMF6^+^ Vγ4 γδ T cells possessed transcriptional profiles consistent with IL-17 and IFN-γ production, respectively.

### Impaired γδ T function in SAP-deficient mice

Finally, we examined the extent to which impaired thymic γδ T cell developmental programming in SAP-deficient mice affected γδ T cell function in peripheral lung and lymphoid organs. A comparison of neonate lung γδ T cells between B6 and B6.SAP^−/−^ mice revealed that there was a significant decrease in lung Vγ4 IL-17, but not IFN-γ, production (**Fig. 6A**) and this decrease corresponded to a significant decrease in SLAMF1^+^IL-17^+^ Vγ4 cells (**Fig. 6B**). No difference was observed in either lung Vγ1 IL-17 or IFN-γ production (**Fig. 6A**), nor did we observe any SAP-dependent decrease in Vγ6 IL-17 production (data not shown). Interestingly, the SAP-dependent decrease in Vγ4 T cell IL-17 production did not persist in the adult lung. Instead, a comparison of adult lung γδ T cell cytokine production between B6 and B6.SAP^−/−^ mice revealed a significant decrease in both lung Vγ1 and Vγ4 IFN-γ production (**Fig. 6C**). We again noted no SAP-dependent decrease in Vγ6 IL-17 production (data not shown). A comparison of adult spleen Vγ1 and Vγ4 cytokine production between B6 and B6.SAP^−/−^ mice also revealed significantly less IFN-γ in SAP-deficient Vγ1 and Vγ4 T cells than in their B6 counterparts, indicating that the SAP-dependent decrease was not tissue-specific (**Fig. 6D**). Together, these data demonstrated that γδ T cell IL-17 and IFN-γ production in the periphery was significantly impaired in SAP-deficient mice. In addition, the data indicated that the SAP-dependent impairment in γδ T cell function in the periphery was both age and subset-dependent.

**Fig. 6.**
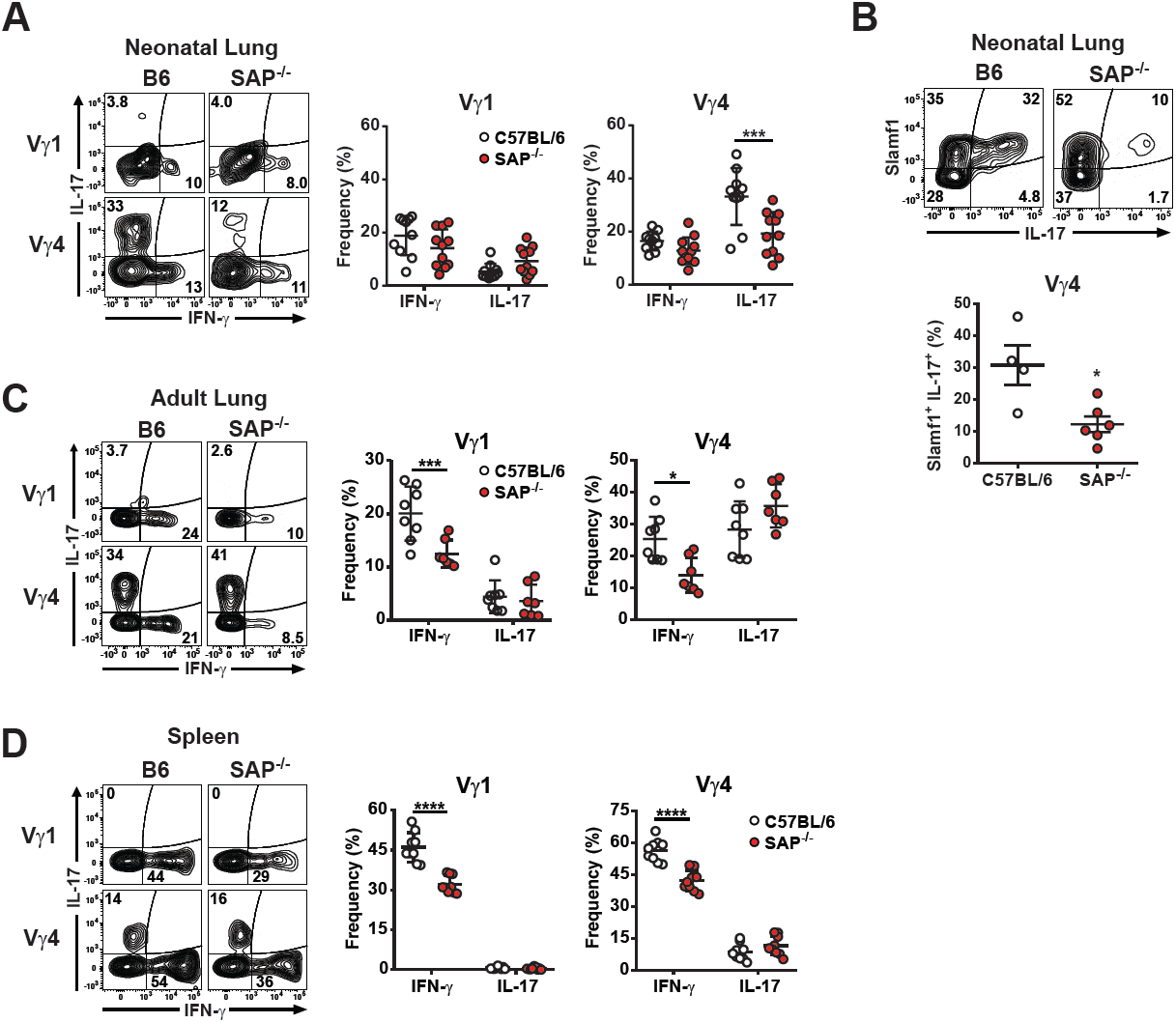
Impaired γδ T cell function in the peripheral tissues in SAP-deficient mice. A) Decreased lung Vγ4 IL-17 production in B6.SAP^−/−^ neonates. Representative contour plots of IL-17 and IFN-γ production from neonate lung Vγ1 and Vγ4 T cells in B6 and B6.SAP^−/−^ mice are shown above. Numbers indicate the percentage of IL-17 and IFN-γ production in each subset. Cumulative frequencies of IL-17 and IFN-γ-producing cells from γδ T cell subsets are shown below. Data are the combined results of 2 independent experiments, day 6 to 7 after birth, sex was not determined. ***p < 0.001 as determined by two-way ANOVA followed by Sidak’s multiple comparisons test. B) Decreased SLAMF1^+^IL-17^+^ lung Vγ4 in neonate day 7 B6.SAP^−/−^ mice. Representative contour plots of SLAMF1^+^IL-17^+^ lung Vγ4 T cells are shown at left. The cumulative data are shown at right, *p < 0.05 as determined by an unpaired, two-tailed t-test. The sex of the mice was not determined. C) Decreased Vγ1 and Vγ4 IFN-γ production in adult B6.SAP^−/−^ lung (C) and spleen (D). Representative contour plots of IL-17 and IFN-γ production are shown at left. Frequencies of IL-17 and IFN-γ-producing cells are shown at right. Data representative the combined results of 2 to 3 independent experiments with male and female mice, 10 to 12 weeks of age, ***p < 0.001, ****p < 0.0001 as determined by two-way ANOVA followed by Sidak’s multiple comparisons test.

## Discussion

Here, we identified the SLAM/SAP signaling pathway as a critical regulator of the functional programming that generates IL-17 and IFN-γ-producing γδ T cells during thymic development. We found that maturation of thymic Vγ1 and Vγ4, was associated with a transition from a SLAMF1SLAMF6 DP phenotype to either SLAMF1 or SLAMF6 SP or SLAMF1SLAMF6 DN phenotypes. This transition was concomitant with the acquisition of γδ T cell effector function in the thymus and in the periphery, where SLAMF1 and SLAMF6 SP phenotypes distinguished IL-17 and IFN-γ-producing Vγ1, Vγ4, and Vγ6 T cells. SLAM family receptor signaling through SAP was critical in this process as SAP-deficient Vγ1 and Vγ4 T cells displayed impaired maturation in the thymus, and impaired effector function in both the thymus and the periphery. Taken together, these findings suggest that the SAP-dependent signaling through SLAM receptors on thymic γδ T cells represents a critical pathway in the developmental programming of γδ T cell effector function.

A number of studies support a model in which high-affinity binding of γδ TCR during development promotes differentiation toward IFN-γ producing cells, while low-affinity interactions favor programming for IL-17 production (22–26, 47). In light of our findings indicating that both IL-17 and IFN-γ production are altered in SAP-deficient mice, it is interesting to note that SLAM receptor signaling has previously been shown to modulate TCR signaling thresholds in conventional αβ T cells (48). In addition, recent reports indicate that the NKT cell differentiation is regulated by TCR signal strength (49, 50), and that the net effect of SLAM receptor signaling is a reduction in NKT TCR signal strength (51). It is possible, therefore, that SLAM/SAP signaling influences γδ T cell developmental programming through the modulation of TCR signals.

The exact developmental stage at which TCR signaling exerts its effect on γδ T cell programming remains an open question. Although it has previously been shown that the CD24^−^ CD44^−^CD45RB^−^ γδ T cell population can give rise to both IFN-γ and IL-17 producers (24), it is unclear whether this population is composed of uncommitted precursors or a mixture of already committed γδ T cells consisting of several functional subsets. Our findings that immature CD24^hi^ thymic γδ T cells are SLAMF1SLAMF6 DP and that mature CD24^lo^ γδ T cells are enriched in SLAMF1 and SLAMF6 SP cells suggest that the CD24^−^CD44^−^CD45RB^−^ γδ T cell population is likely a heterogeneous mixture of SLAMF1 and SLAMF6 SP γδ T cells. The decrease in CD24^lo^ γδ T cells observed in SAP-deficient mice coupled with the loss of SLAMF1 SP γδ T cells is consistent with a role for SAP in both the maturation and acquisition of function. Because the co-expression of both SLAMF1 and SLAMF6 on immature CD24^hi^ γδ T cells puts them in an optimal position to influence TCR signal strength, it could be possible that SLAM/SAP signaling modulates the TCR signals received by the immature γδ T cell, thereby influencing its differentiation. An important caveat to this model is that the frequency and function of Vγ6 T cells, whose development has been demonstrated to be influenced by TCR signal strength (25), did not appear to be dependent on SAP. Whether the effect of SAP is due to its effect on TCR signaling, or is due to a TCR-independent process will require additional investigation.

Increasing evidence suggests that the development of IL-17- and IFN-γ-producing γδ T cells is also regulated by TCR-independent mechanisms. γδ T cell development is dependent on a characteristic subset of transcription factors including SOX4, SOX13, Blk, c-maf, HEB, and PLZF, among others (27, 28, 45, 52–54). Recently, it was reported that at least some γδT17 programming results from a SOX13-dominated transcriptional hard-wiring of early thymocyte precursors (29). These SOX13 progenitors appear to give rise to IL-17-producing Vγ4, but not Vγ6 cells. Our observation that Vγ4, but not Vγ6 subsets are SAP-dependent are therefore consistent with a role for SLAM receptors and SAP in the initiation of a transcriptional program critical for the acquisition of function. Indeed, because SLAM receptors are expressed on hematopoietic stem cells (55–57), they are in a position to influence γδ T cell programming at the earliest stages of development.

Our findings demonstrate the presence of distinct SLAM family receptor expression profiles on γδ T cell subsets in the periphery that are tightly associated with function. These findings not only confirm a previous report that SLAMF1 marks γδT17 cells (42), they reveal the presence of a mutually exclusive expression pattern of SLAMF1 and SLAMF6 on γδ T cells that is tightly associated with distinct transcriptional programs and with IL-17 and IFN-γ production, respectively. The finding that SLAM family receptor expression patterns are established during thymic development, and that they are already associated with function as early as embryonic day 17, suggests that they result from a developmental program.

In support of this notion, significant changes in SLAMF1, SLAMF4, SLAMF6, and SLAMF7 expression have been described on B cells as they progress through developmental stages (58), and SLAM receptor expression profiles on hematopoietic stem cell and multipotent progenitor cell populations (55) have recently been demonstrated to mark functionally distinct progenitor populations (57). Together, our findings suggest the possibility that the coordinate signaling of distinct arrays of SLAM family receptors plays a role in the development of γδ T cells in the thymus. This would be consistent with reports indicating that multiple SLAM receptors are involved in NKT cell development (34, 51, 59, 60), and may explain the inability to observe γδT17 defects in mice lacking only a single SLAM family receptor (42).

Our observations indicated that SAP deletion did not completely abolish the development of IL-17-producing Vγ4, but instead reduced their numbers in thymus and neonatal lung by approximately 40%. This finding likely suggests the existence of one or more specific SAP-dependent Vγ4 subsets, as is the case with the Vγ1 T cell population that is composed of the SAP-dependent Vγ1Vδ6.3 subset and other SAP-independent subsets. Recent data demonstrating that the TCR repertoire of IL-17-producing Vγ4 T cells is highly restricted (61, 62), suggests the possibility that the partial decrease in Vγ4 IL-17 production observed here may be due to a complete failure in the development of a specific IL-17-producing Vγ4 subset. Such a model is also consistent with the observation that the SAP-dependent decrease in lung γδ T cell IL-17 production observed in neonates was not consistently observed in adult mice. We speculate that the restricted development of IL-17-producing Vγ4 T subsets to embryonic and neonatal thymus (21) results in decreased SAP-dependent Vγ4 T subsets in the neonatal lung, and that over time, homeostatic proliferation of SAP-independent Vγ4 T subsets results in a repopulation of the adult lung. A future analysis of the γδ TCR repertoire in SAP-deficient mice should shed light on this issue.

In summary, our findings demonstrate that the SLAM/SAP signaling pathway plays a key role in shaping the balance of IL-17- and IFN-γ producing γδ T cell commitment during thymic development. Future studies aimed at understanding the specific mechanisms of SLAM family receptor signaling in this process will be required to integrate this pathway into existing models, and should provide insight into the central mechanisms that determine γδ T cell functional programming.

## Acknowledgements

We wish to thank Roxana del Rio Guerra and the Flow Cytometry Cell Sorting Facility for help in cell sorting, Jessica Hoffman and Scott Tighe and the Vermont Integrated Genome Resource for help in microarray and qPCR, Wendy Havran for providing a protocol for skin γδ T cell isolation, Cynthia Baldwin and Janice Telfer for helpful discussion, and Willi Born and Rebecca O’Brien for the 17D1 hybridoma.

## Supplementary Figure legends

**Fig. S1.**
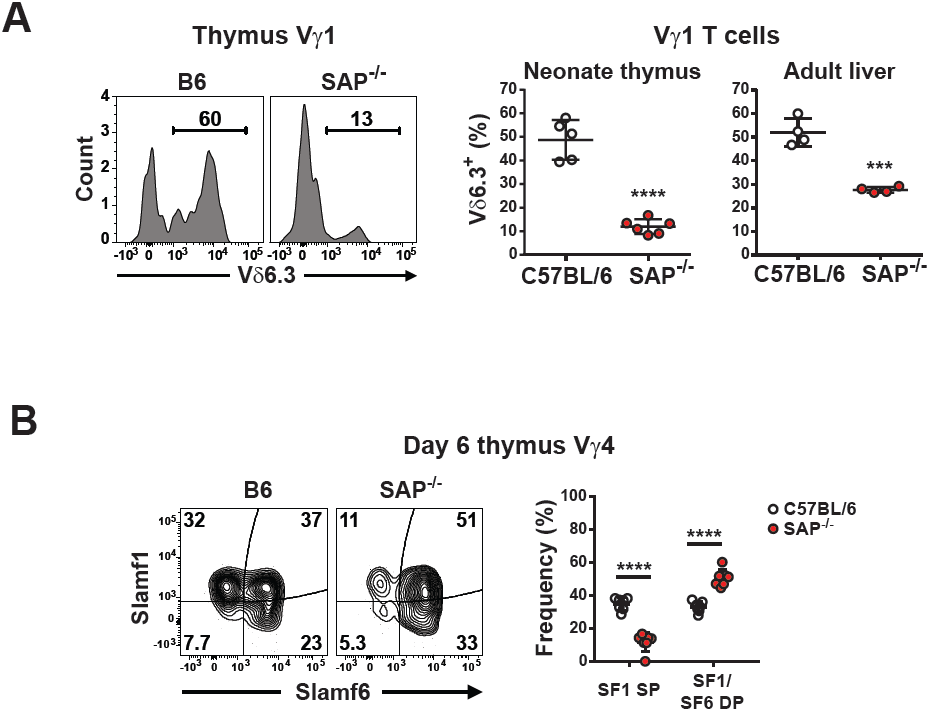
SAP-dependent development of γδ T cell subsets. A) SAP-dependent development of Vγ1Vδ6.3 T cells. Representative histograms of Vδ6.3 expression on neonatal thymic Vγ1 T cells are shown at left, and cumulative frequencies for day 5 neonatal thymus and adult liver (8.5 weeks of age males) are shown at right, ***p < 0.001, ****p < 0.0001 as determined by an unpaired, two-tailed t-test. Data are representative of 2 independent experiments, 4-6 mice per group. B) Comparison of SLAMF1 SP and SLAMF1SLAMF6 DP Vγ4 T cells from day 6 thymus between B6 and B6.SAP^−/−^ mice. Representative contour plots are shown at left and cumulative data are shown at right, ****p < 0.0001.

**Fig. S2.**
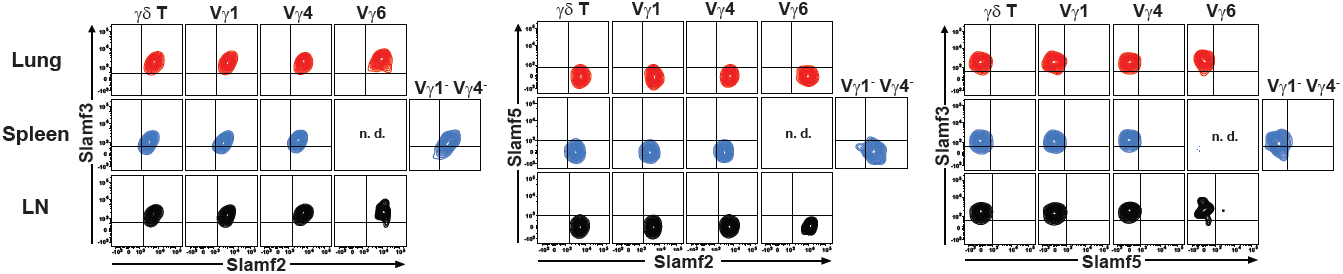
Expression of SLAMF2, SLAMF3, and SLAMF5 on γδ T cell subsets from C57BL/6J lung, spleen, and lymph node (LN). Representative contour plots are shown. Data are representative of 2 separate experiments, 4 female mice per group, 8 weeks of age, n.d., not detected.

## Notes

1 This work was supported by NIHR21AI119974, NIHP30GM118228, and NIHS10OD018175 (JEB).

